# Comparative transcriptomic analysis reveals key components controlling spathe color in *Anthurium andraeanum* (Hort.)

**DOI:** 10.1101/2020.12.29.424726

**Authors:** Jaime A. Osorio-Guarín, David Gopaulchan, Corey Quackenbush, Adrian M. Lennon, Pathmanathan Umaharan, Omar E. Cornejo

**Author notes:** These authors contributed equally to this work (Jaime A. Osorio-Guarín, David Gopaulchan).

## Abstract

*Anthurium andraeanum* (Hort.) is an important ornamental in the tropical cut-flower industry. However, there is currently not enough information to establish a clear connection between the genetic model(s) proposed and the putative genes involved in the differentiation between colors. In this study, 18 cDNA libraries related to the spathe color and developmental stages of *A. andraeanum* cut-flowers were characterized by transcriptome sequencing technology. For the *de novo* transcriptome, a total of 114,334,082 primary sequence reads were obtained from the Illumina sequencer and were assembled into 151,652 unigenes. Approximately 58,476 transcripts were generated and used for comparative transcriptome analysis between three varieties that differ in spathe color (‘Sasha’ (white), ‘Honduras’ (red), and ‘Rapido’ (purple)). A large number of differentially expressed genes (8,324) that were potentially involved in multiple biological and metabolic pathways were identified, including the flavonoid and anthocyanin biosynthetic pathways. Our results showed that chalcone synthase (*CHS*) and flavonoid 3’-hydroxylase (*F3’H*) were the main genes differentially expressed in the white/red/purple comparison. We also identified a differentially expressed cytochrome *P450* in the late developmental stage of the purple spathe that appeared to determine the difference between the red- and purple-colored spathes. Additionally, putative MYB-domain protein candidates that could be responsible for the control of the biosynthetic pathway were identified. The results provided basic sequence information for future research on spathe color, which have important implications for breeding strategies in this ornamental.

**Core ideas:**

- RNA-seq was performed on three anthurium varieties.
- Gene expression was compared for developmental stage and spathe color.
- Differentially expressed unigenes were identified.
- Putative MYB-domain protein candidates of the anthocyanin biosynthetic pathway were identified.

## INTRODUCTION

The large diversity of floral traits in angiosperms have fascinated researchers and ornamental flower enthusiasts alike for generations. Evolutionary biologists have long held the view that the large diversity of floral traits is associated with high rates of diversification among species; a process that is likely mediated by natural selection (Sobral et al., 2015). Among floral traits, flower color and scent are undoubtedly of great evolutionary importance given the direct impact that they can have on pollinator choice and thus fitness of the plant (Jones and Reithel, 2001; Hopkins and Rausher, 2012; Hirota et al., 2013).

*Anthurium andraeanum* (Hort.) is a species complex created by interspecific hybridization between *A. andraeanum* Linden ex André and other related species (Kamemoto and Kuehnle, 1996). This species has a brightly coloured spathe surrounding the inflorescence and its color is one of the most attractive characteristics that gives the species commercial value. The large diversity in spathe colors such as, white, red, pink, orange, coral, green, brown and purple and patterns such as, obake, striped and colored borders (Kamemoto et al., 1988; Wannakrairoj and Kamemoto, 1990; Kamemoto and Kuehnle, 1996) has made this species an important tropical ornamental crop (Dufour and Guérin, 2003). In addition, it is well known that anthocyanins (synthesized in the cytosol and localized in vacuoles), widely found in the flowers, seeds, fruits and vegetative tissues of vascular plants as soluble flavonoid pigments, also participate in defense against a variety of biotic and abiotic stressors in plants (Tanaka et al., 2008; Santos-Buelga et al., 2014; Khoo et al., 2017). Besides being directly beneficial to plants, anthocyanins have also been used as natural food colorants and display vital nutraceutical properties that could be advantageous for human health (Tanaka et al., 2008; Pojer et al., 2013; Li et al., 2017).

Extensive crossing analyses have been done on varieties of *Anthurium* to unveil the genetic mechanisms involved in color formation and segregation (Kamemoto et al., 1988; Elibox and Umaharan, 2008). Elibox and Umaharan (2008) showed that spathe color is controlled by duplicate recessive epistasis and proposed two genes (O and R) as responsible for the distinction between color vs white and a modifier gene (M) as responsible for the distinction between reds and pinks/oranges (Elibox and Umaharan, 2008).

The biosynthetic pathway generating anthocyanins is highly conserved in plants and the genes involve has been extensively studied (Holton and Cornish, 1995). For example, the chalcone synthase (*CHS*) catalyzes the first reaction leading to anthocyanin biosynthesis and assists in forming the intermediate chalcone, which is the primary precursor for all classes of flavonoids. Dihydroflavonol 4-reductase (*DFR*) is the first committed enzyme of anthocyanin biosynthesis in the flavonoid biosynthetic pathway and is responsible for the formation of leucoanthocyanidins which can be converted into colored anthocyanidins by anthocyanidin synthase (*ANS*) (Tanaka et al., 2008). The enzymes which catalyze specific steps of the anthocyanin biosynthesis pathway are encoded by structural genes which are in-turn under the control of regulatory genes (transcription factors). For example, transcription factors like R2R3-MYB and members of the basic helix-loop-helix (bHLH) (R/B) family, have been reported as regulatory elements that controls pigmentation in flowers and fruits in different species (Springob et al., 2003; Quattrocchio et al., 2006; Lin-wang et al., 2010; Motamayor et al., 2013; Singh et al., 2014). A putative transcription factor AaMYB2, ectopically expressed in tobacco increased anthocyanin accumulation and increased the expression of *DFR*, flavonoid 3’-hydroxylase (*F3’H*), *ANS*, and possibly *CHS* genes (Li et al., 2016). However, regulatory genes driving the expression of the enzymes in the pathway might differ depending on the species (Mori et al., 2009; Lin-wang et al., 2010; Motamayor et al., 2013; Tan et al., 2013; Lou et al., 2014; Singh et al., 2014; Cao et al., 2015; Zhao and Tao, 2015; Li et al., 2016).

In *A. andraeanum*, the major color pigments in the spathe are anthocyanins, predominantly cyanidin and pelargonidin derivatives, of which the content and ratio determine the color and its intensity (Williams et al., 1981). A study showed that the interaction between a R2R3-MYB transcription factor with a basic helix-loop-helix (AabHLH1) regulates the proanthocyanidin accumulation (Li et al., 2019). Besides, temporal and spatial expression analysis of four genes involved in flavonoid synthesis (*CHS, F3’H, DFR* and *ANS*) suggests that *DFR* transcript levels vary significantly between developmental stages and might represent a relevant regulator (Collette et al., 2004). A phenotype and transcriptome analysis of anthurium leaf color mutants suggested that color formation in leaves was greatly affected by chloroplast development and pigment biosynthesis with increased expression of flavonoid 3’5’ hydroxylase (*F3’5’H*) and *DFR* in the dark green leaf mutant compared to the wildtype (Yang et al., 2015). However, the genes involved in the regulation of color in these mutants could not be identified. Thus, a more detailed analysis of gene expression was essential to improve our understanding between the genetic model(s) and the putative regulatory genes involved in the differentiation between colors.

Approaches such as RNA sequencing (RNA-Seq) have allowed the profiling of global gene expression patterns and the mapping of simple or complex traits, respectively, both in model and non-model plants. In this work, we performed a comprehensive transcriptomic analysis of white, red, and purple spathes of three varieties of *A. andraeanum* from two cut-flower development stages. The aims of our study were: 1) to identify differentially expressed genes directly involved in the biosynthesis of anthocyanins, and 2) to identify putative regulatory genes involved in the generation of spathe color. The present study provides an important molecular basis for understanding the color formation mechanism for the germplasm evaluation and ornamental breeding.

## MATERIALS AND METHODS

### Plant material

*A. andraeanum* cut-flowers were collected between 6 am and 8 am from mature 3-to 4-year-old plants grown in shade houses with 75% shade and approximately 12 h day length at Kairi Blooms Ltd., a commercial anthurium farm located at Carapo village, Trinidad.

To identify the genes differentially expressed between white (W), red (R) and purple (P) spathes, cut-flowers were collected from ‘Sasha’, ‘Honduras’ and ‘Rapido’ cultivars respectively (Figure 1). Samples were harvested in triplicate (3 individual spathes from different plants) from two stages of cut-flower development: early (E) (stage 2 -cut-flower first visible) and late (L) (stage 6-spathe newly opened and fully expanded) (Figure 1). All samples were harvested and immediately stored in liquid nitrogen.

**Figure 1.**
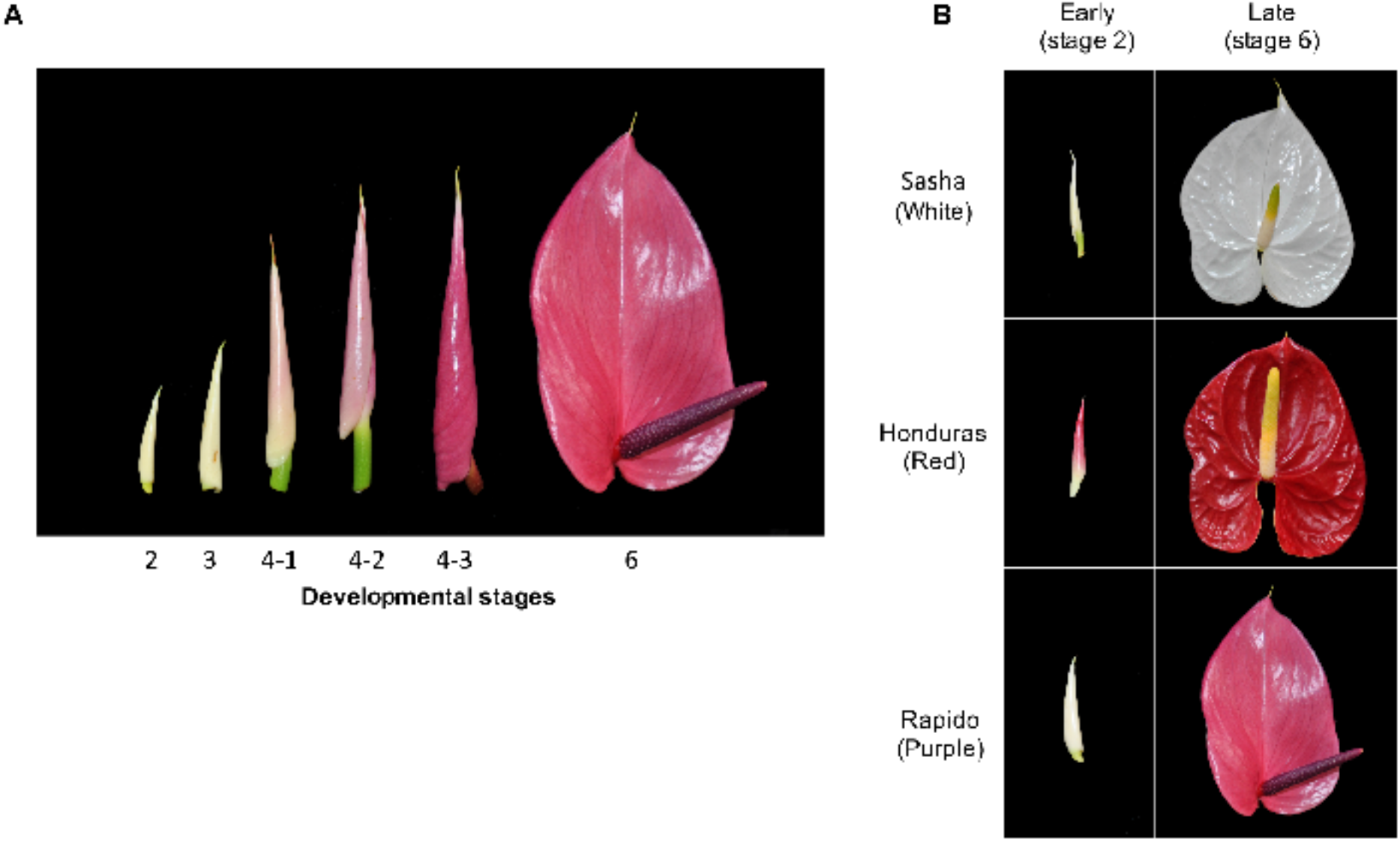
A. Stages of spathe development for *Anthurium andraeanum*, B. Stages 2 and 6 used for each of the three varieties for the RNA-seq assay.

### RNA isolation, library preparation and sequencing

The total RNA was isolated from the spathe tissue (80 mg) samples (three varieties x two development stages x 3 biological replicates) using the RNAqueous Kit containing the Plant RNA Isolation Aid (Applied Biosystems, Foster City, CA) as described by the manufacturer. The RNA was treated with Turbo DNA-free (Applied Biosystems, Foster City, CA) and 0.1 volume of 3 M sodium acetate and 2.5 volumes of 100% ethanol were added to each sample. Samples were then shipped in dried ice to Washington State University where they were purified with ethanol (70%) and resuspended in 25 µL of ultrapure water. A Qubit fluorometer was used to quantify the amount of total RNA and the quality of each sample was determined on an Agilent 2100 Bioanalyzer. All samples were normalized so that each contained 1 µg of total RNA in a volume of 10 µL.

To construct the RNA libraries, the Illumina TruSeq Stranded Total RNA with Ribo-Zero Plant kit were used according to the manufacturer’s instructions. To assess the quality and determine the molarity of the prepared libraries, each sample was run on an Agilent 2100 Bioanalyzer. The samples were then normalized to 10 nM, randomized, and sequenced in a single lane (for a total of two lanes) of Illumina HiSeq 2500 System (paired-end, 100 bp reads) at the Washington State University Genomics Core.

### *De novo* transcriptome of anthurium and functional annotation

The reference transcriptome was derived from the spathes of ‘Honduras’ cut-flowers. Samples were harvested in triplicate (3 individual spathes from different plants) from spathe developmental stages 2, 4-1 (newly extended peduncle with a peduncle length of 3–4 cm) and 6. Total RNA was extracted as previously described and combined into one pool and shipped to Beijing Genomics Institute (BGI), Shenzhen, China to develop a *de novo* reference transcriptome. RNA quality was verified using an Agilent 2100 Bioanalyzer (Agilent Technologies, Inc., Santa Clara, CA) and poly (A) mRNA was isolated with oligo (dT) magnetic beads. The poly (A) mRNA was fragmented into 200 to 700 bp and first-strand cDNA was synthesized with random hexamer primers followed by the synthesis of the second strand with buffer, dNTPs, RNase H and DNA polymerase I. The double strand cDNA was purified and subjected to end repair and base A addition. Fragments were ligated with sequencing adapters, then purified by agarose gel electrophoresis (200 bp size selection) and enriched by PCR amplification. Primary sequence data (paired-end, 91 bp reads) was generated using the Illumina HiSeq 2000 (Illumina, Inc., San Diego, CA). We used the Trinity software (Grabherr et al., 2011) with default parameters, to assemble reads. The software’s TGICL (Pertea et al., 2003) and Phrap (Vogel et al., 2006) were utilized to get sequences that could not be extended on either end.

Unigenes were aligned with blastx (expect value < 10^−5^) to NCBI non-redundant protein (Nr), Swiss-Prot, Kyoto Encyclopedia of Genes and Genomes (KEGG) and Cluster of Orthologous Groups (COG) databases, to identify proteins with the highest sequence similarity to the respective unigene along with their functional annotation. Proteins with highest ranks in the BLAST results were used to determine the coding regions in the unigenes and translate them into peptide sequences. To further characterize the function of the genes identified, DEGs and consensus sequences of isoforms were mapped against the UniProtKB/Swiss-Prot database using an expect value threshold of 1×10^−5^. The BlastX output was subjected to Blast2GO for Gene Ontology (GO) analysis.

### Analysis of RNA sequencing data

Once reads were obtained, samples were demultiplexed using the standard Illumina pipeline. Quality per sample was assessed using FastQC v0.11.1 (Andrews et al., 2012). Individual samples were quality and adaptor trimmed using Trim Galore v0.5.0 and Cutadapt v2.10 (Martin, 2011; Krueger, 2018). For the quality trimming, we set sequences to be trimmed for bases with Illumina Phred scores lower than 25 and sequences with less than 50% of the original length of the read were removed from the dataset. For adaptor trimming we considered an error rate of 0.2.

Mapping and alignment of reads to transcripts was performed with bowtie2 v2.4.2 (Langmead et al., 2009; Langmead and Salzberg, 2012), keeping all ambiguously mapping positions. All mapping positions were maintained to estimate, using a probabilistic approach, the number of reads supporting the expression of any given transcript, as implemented in RSEM (Haas et al., 2013). We fitted a generalized linear model with color and stage treatments with a negative binomial using the quasi-likelihood approach to reduce the impact of false positives, as recommended by edgeR developers (McCarthy et al., 2012) and followed the experimental design depicted in Figure 1B. In this model, we tested simultaneously for the effect of developmental stage (E and L) and color (W, R, and P) and controlled for differences in library size. We set two sets of contrasts *a priori*, in which we aimed to identify differential expression between colors at each stage (Figure 2).

**Figure 2.**
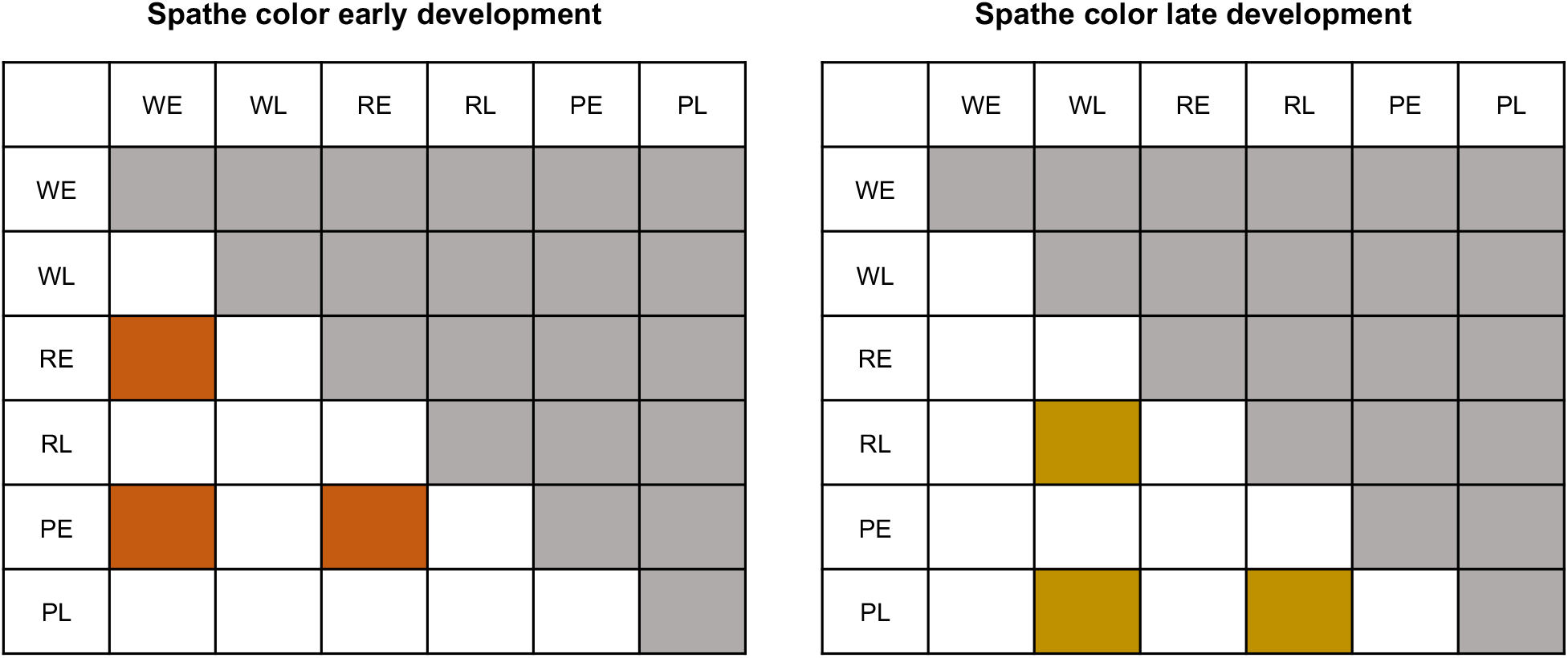
Matrices depicting specific contrasts designed to identify genes differentially expressed. Comparisons between spathes of different colors (W, R,P) in stage 2 (Color Early (E)); and between spathes of different color in stage 6 (Color Late (L)).

A multidimensional scaling (MDS) was used to visualize the overall variation in the samples. Normalized read counts from the top 10,000 variable genes were analyzed with MDS first across all samples and then within each treatment separately using plotMDS in the limma package (Ritchie et al., 2015) in R (R development core team, 2008). An initial screening of the data was carried out through a hierarchical clustering dendrogram for each treatment based on the Euclidian distance and grouped with the unweighted pair group method with arithmetic mean (UPGMA) using the dendrogram function of the ape package (Paradis et al., 2004) of R.

Differentially expressed genes (DEGs) were calculated by using the total number of transcripts per million (TPM) for filtering purposes and using the estimated effective counts of reads mapping to each transcript with the software package edgeR (Robinson et al., 2010; McCarthy et al., 2012). DEGs were defined as genes having a false discovery rate (FDR) (Benjamini and Hochberg, 1995) < 5×10^-4^ and an absolute log fold change (logFC) ≥ 2.

Finally, for the genes that present a function of interest, we took the Log_2_ of the TPM and estimated a centered and normalized measure of expression in the following way:

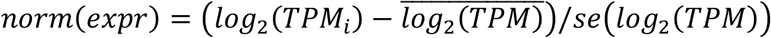

where *i* corresponds to individual measure of expression for that gene in individual sample *i*. All differences in expression for individual genes of interest are shown for these individual genes in the normalized form. All subsequent statistical analyses and graphical representations were generated in R.

### Identification of putative transcription factors

An expression network was used to identify potential transcription (specifically MYB domain transcription factors) responsible for the regulation of differential expression of genes involved in the anthocyanin metabolism. For this, we performed a weighted correlation network analysis (WGCNA) as implemented in WCGNA package (Langfelder and Horvath, 2008) in R. We manually searched for the best thresholding power conditions for the construction of the network and searched for modules of potentially co-regulated genes in association with differences in color between spathes. We extracted all of the transcripts found in each module and identified the modules containing genes involved in anthocyanin metabolism that showed significant differential expression (p ≤ 0.0005). Then, we extracted the putative MYB domain containing genes that were identified as being in the same module that contained differentially expressed genes of interest (involved in anthocyanin metabolism) and retained those with the highest correlation in expression.

## RESULTS

### *De novo* transcriptome and RNA sequencing

For the *de novo* transcriptome, a total of 114,334,082 primary sequence reads were obtained from the Illumina sequencer. After stringent quality control checks and filtering, 105,143,382 high-quality clean reads (9,462,904,380 nucleotides) were identified. Among the reads, 98.08% of the nucleotides had quality values > Q20, while the proportion of unknown nucleotides was < 0.00% and the GC content was 42.97%. Filtered reads were assembled into 151,652 unigenes, the presence of unknown bases in the sequences varying between 0% to 5% and the length of the unigenes ranged from 201 to 12548 bp.

For the RNAseq, we obtained a total of 391.29 million clusters (782.58 million reads) of paired-end sequencing with Illumina Phred qualities ≥ Q30. The expected number of reads per sample was 21.78 million reads or 5.5% of the total reads. The distribution of the reads among samples was relatively even across samples, with samples having between 3.8% and 6.7% of the total reads (Supplemental Figure S1). On average, 85% of reads mapped to the reference transcriptome per sample. There was a total of 151,652 transcripts in the reference transcriptome. There were 58,476 transcripts for which the number of mapped reads across samples exceeded a minimum threshold of TPM on average across samples. One of the samples (ER2) was excluded from the analysis because it showed a very low rate of mapping to the reference transcriptome and was a clear outlier based on the UPGMA cluster analysis (Supplemental Figure S2). Finally, the MDS analysis clearly separated the stages in principal component 1 and spathe colors in principal component 2 (Figure 3).

**Figure 3.**
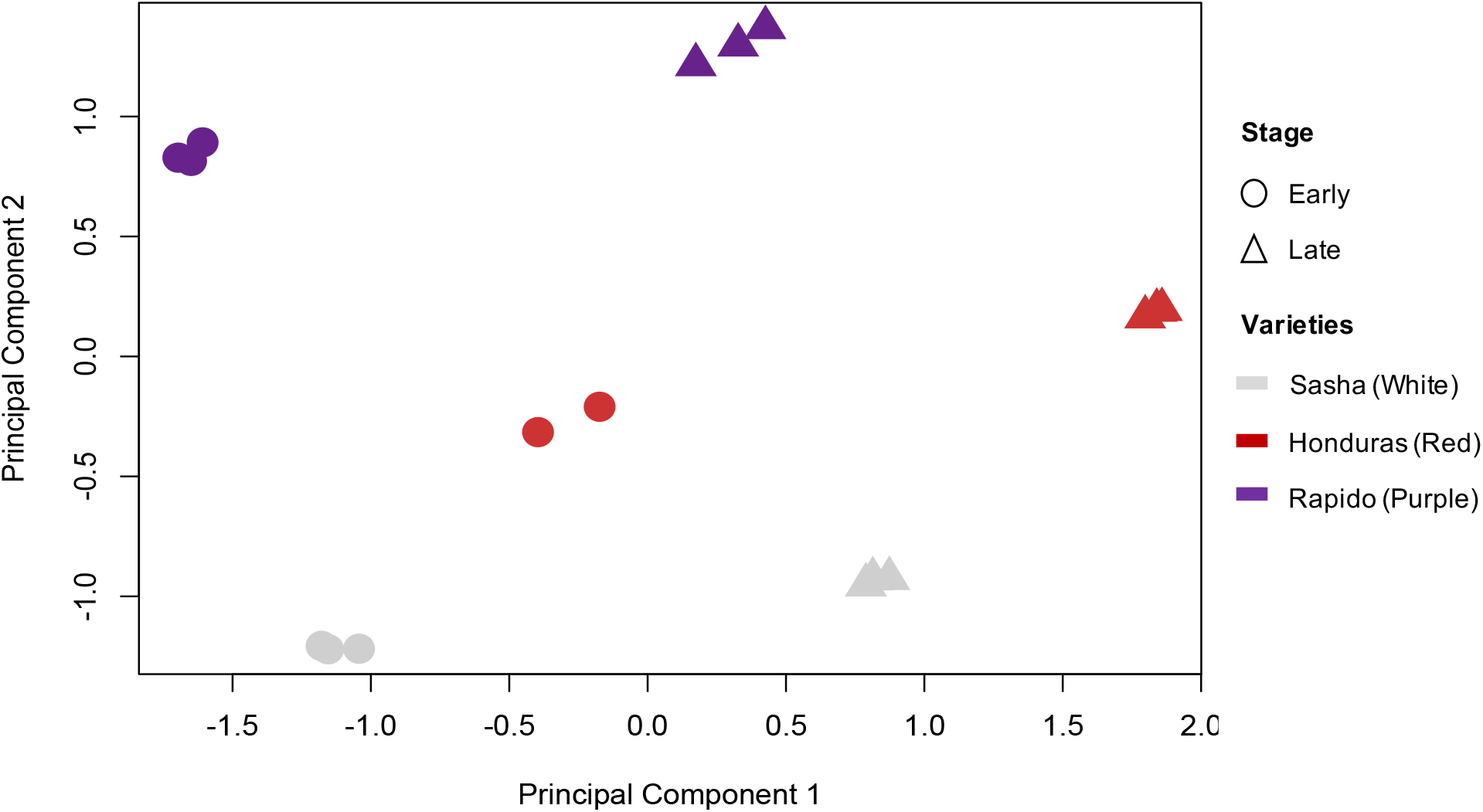
Multidimensional scaling based on gene expression. Samples on the negative scale of principal component 1 correspond to early developmental stages (circles) for either color, and samples on the positive side correspond to late developmental stages (triangles). Dispersion along principal component 2 explained variation between different colors: white (light grey), red (red) and purple (purple).

### Differential expression between stages and across different color spathes

We identified 8,324 differentially expressed transcripts at a conservative FDR threshold value of 5×10^−4^ and logFC ≥ 2 for all the pairwise contrasts after fitting a quasi-likelihood model. There was a similar number of genes differentially expressed between stages within each color of spathe (Table 1 and Figure 4). The relative number of DEGs between colors in the early stage of spathe development was generally smaller than the number of DEGs in late stage. We found groups of genes showing similar patterns of differential expressions across comparisons (Figure 5). Based on annotations, we identified that a large number of genes involved in transcriptional regulation and cell wall expansion were differentially regulated between stages for all colors. The pattern of differential expression was more complex between colors within each developmental stage, which is consistent with the fact that between the varieties, the spathes were also highly differentiated in shape in addition to color. A comprehensive list of differentially expressed genes for each comparison is described in the Supplemental Table S1. Quantitative set diagrams showed how differentially expressed genes were shared across comparisons for each set of contrasts (Figure 5). In our stage comparison (E vs L), we observed for all colors a general trend of having more statistically significant downregulated genes than upregulated in the late stage. Although we did not find a remarkable difference between the number of downregulated and upregulated genes in the rest of the comparisons, the general trend suggested that there was a slightly larger number of downregulated genes in colored spathes (R, P) when compared to the white (W) spathe at both development stages (E, L) (Supplemental Table S1).

**Table 1.**
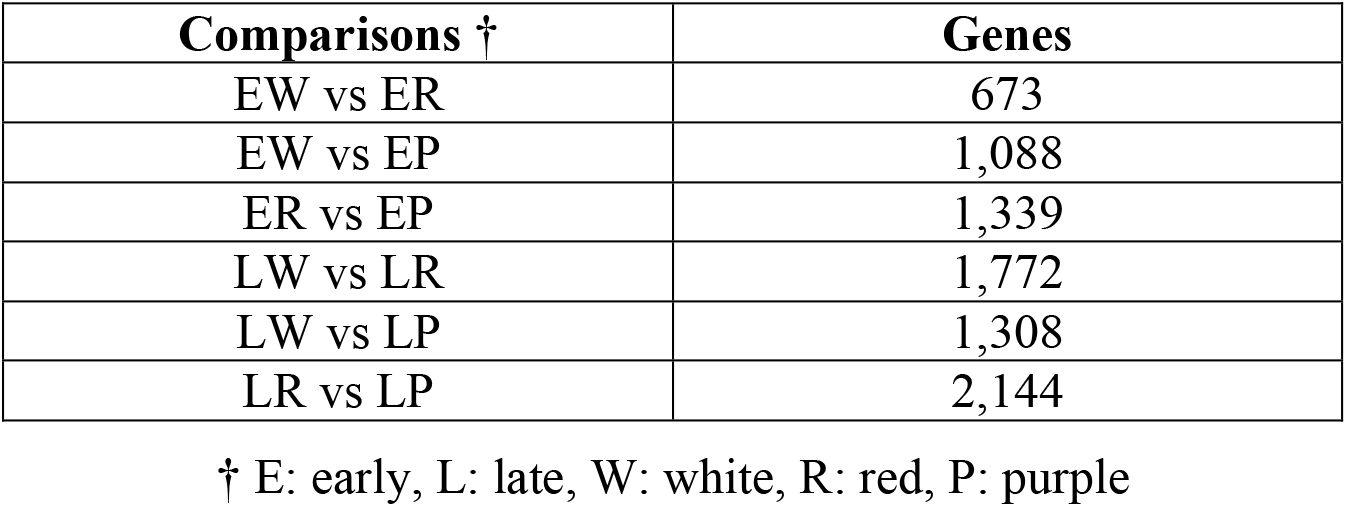
Number of genes differentially expressed between stages and spathe colors.

| Comparisons † | Genes |
| --- | --- |
| EW vs ER | 673 |
| EW vs EP | 1,088 |
| ER vs EP | 1,339 |
| LW vs LR | 1,772 |
| LW vs LP | 1,308 |
| LR vs LP | 2,144 |

† E: early, L: late, W: white, R: red, P: purple

**Figure 4.**
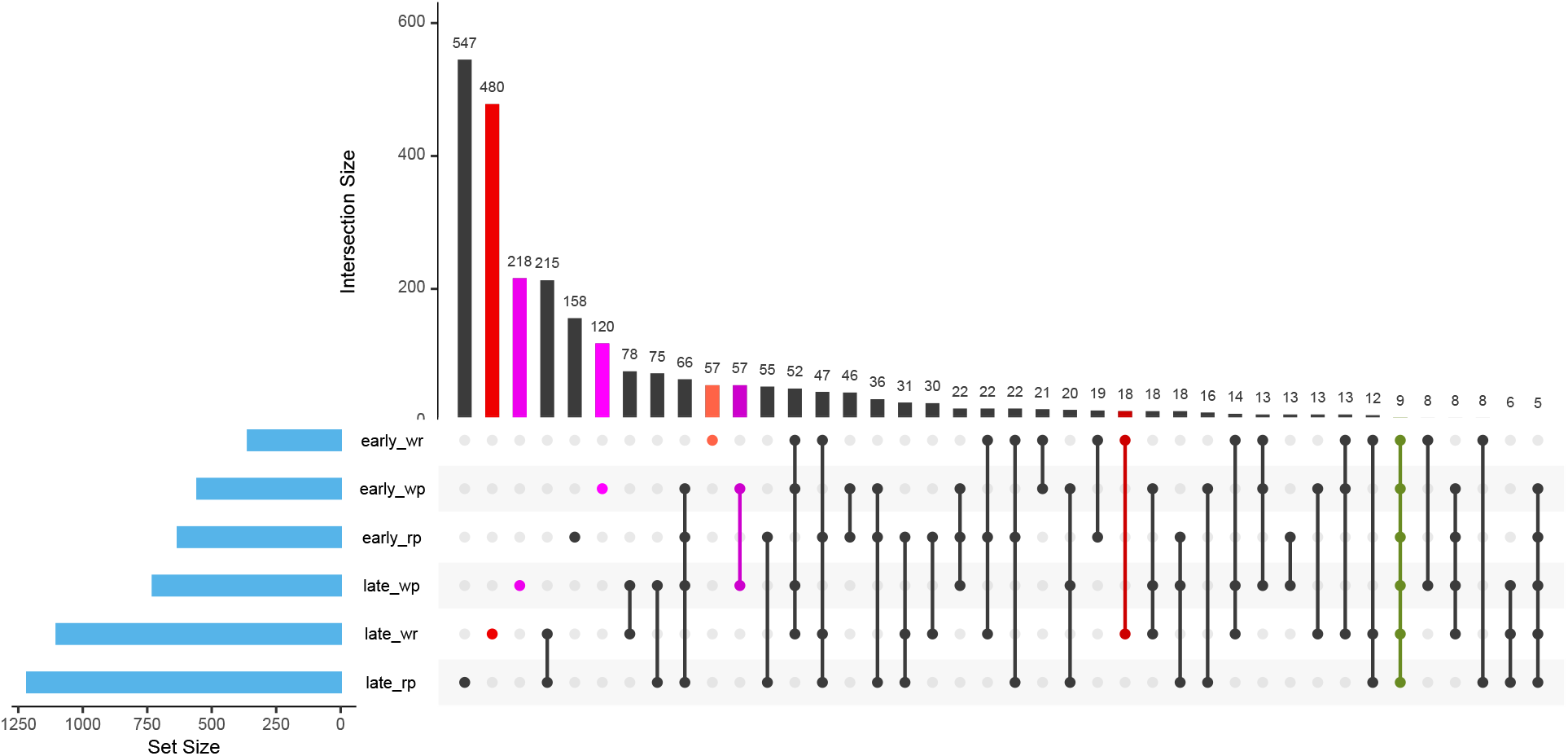
Venn diagram of the differentially expressed genes (DEGs) of RNA-Seq. Venn diagram showing the number of differentially expressed genes (FDR threshold value of 5×10^−4^ and logFC ≥ 2) between comparison of developmental stages and spathe colors.

**Figure 5.**
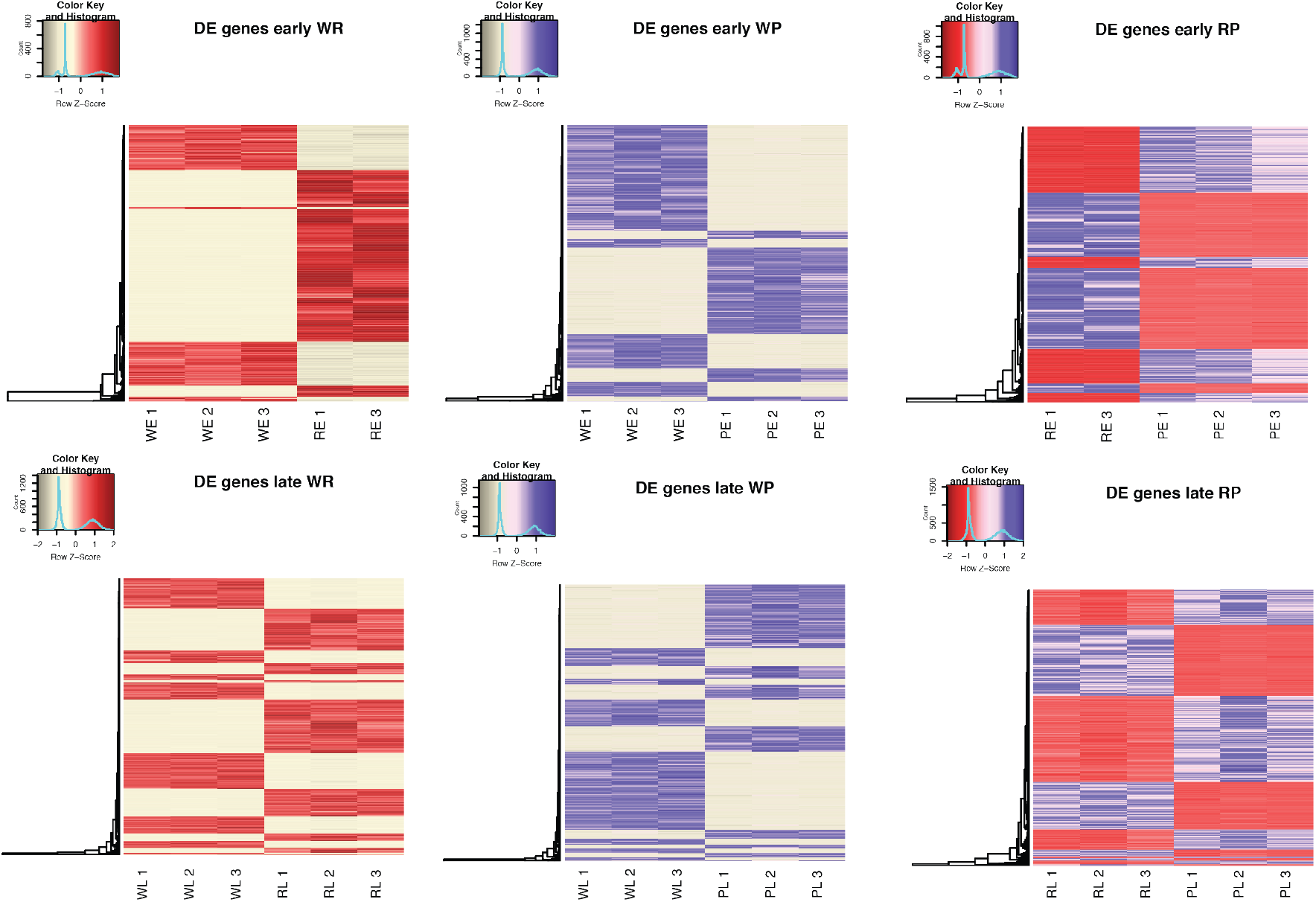
Heat map of RNA-Seq transcriptome analysis for 8,324 selected genes from the *Anturium andraeanum.* For each illustration, W: white, R: red, P: purple; and to stages: E: early, L: late. The biological replicates for each combination of color stage are indicated with numbers (1,2,3). Only genes with log fold change (logFC) ≥ 2 and at an FDR < 5×10^−4^ are represented in the heatmaps. In figures, lighter colors correspond to under-expressed genes and darker colors to over-expressed genes.

We found a large number of DEGs in our comparisons, but we focused our attention to those genes involved in the anthocyanin biosynthetic pathway (Figure 6). We found that the *CHS* gene, at the top of the regulatory cascade of the anthocyanin biosynthetic pathway, had significantly higher expression levels in red and purple spathes than in white spathe in both early and late stages of development (Figure 7A). Additionally, we found that the *F3’H* gene had a significantly higher expression in colored spathes when compared to white at all stages (Figure 7A). Interestingly, we also found a homologous gene to a cytochrome *P450* to be differentially expressed between red and purple flowers at the late stage (Figure 7B).

**Figure 6.**
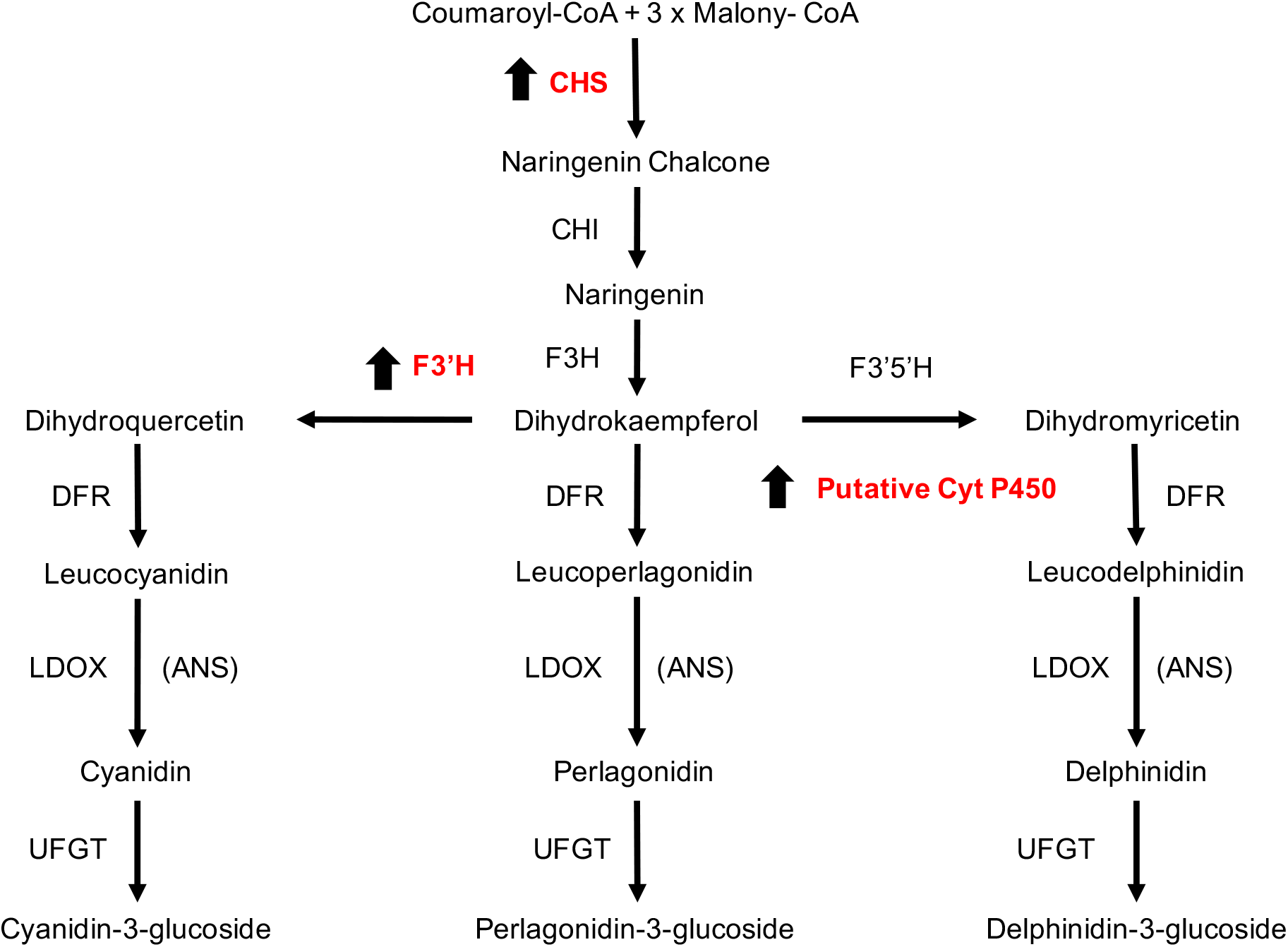
Adapted diagram showing the anthocyanin biosynthetic pathway. The arrows correspond to upregulated genes. CHS: chalcone synthase; CHI: chalcone isomerase; F3H: flavanone 3-hydroxylase; F3’H: flavonoid 3’-hydroxylase; F3’5’H: flavonoid-3’,5’-hydroxylase; DFR: dihydroflavonol 4-reductase; LODX-ANS: anthocyanidin synthase; UFGT: UDP-Glc-flavonoid 3-O-glucosyl transferase; and FLS: flavonol synthase. Modified from Holton and Cornish (1995).

**Figure 7.**
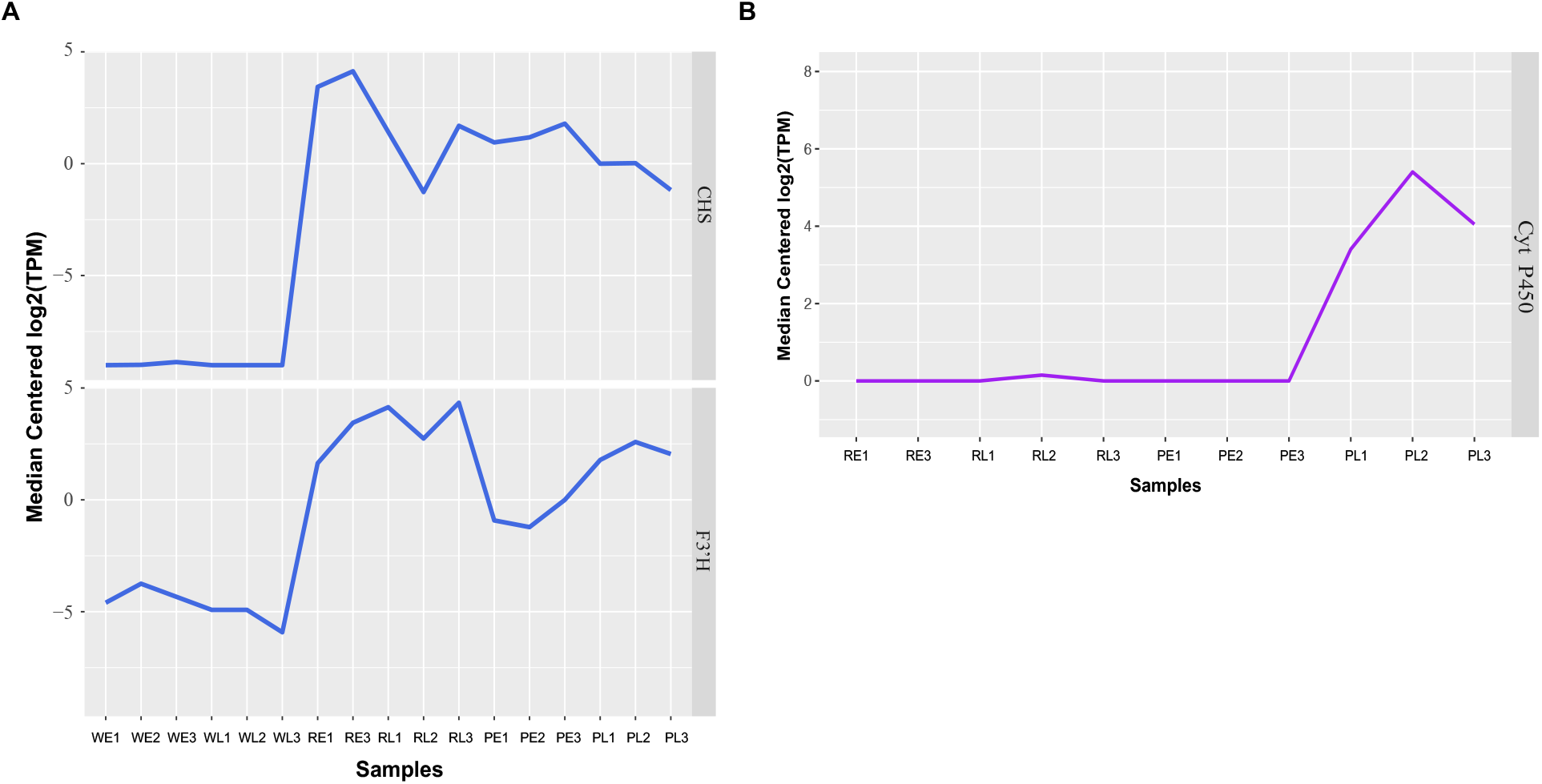
Measure of expression based on Log2 of transcripts per million (TPM). A. The genes *CHS* and *F3’H* were upregulated in red or purple when compared to white and, B. The putative gene *cyt P450* was upregulated in late stages of purple flowers when compared to red.

### Identification of putative transcription factors

It has been shown that *CHS* tissue-specific regulation is controlled by a MYB-like domain protein in different systems (Lin-wang et al., 2010; Motamayor et al., 2013; Singh et al., 2014). The *Anthurium* transcriptome, and many plant genomes have a large number of annotated MYB domain and MYB-like domain proteins (Supplemental Table S2). In order to reduce the number of potential candidates that could be involved in the control of *CHS* differential expression, we performed a WGCNA of expression across all samples (Figure 8A). The general results of the clusters of genes showing similar patterns of expression are shown in Figure 8B. Our analysis suggested that 21 modules or clusters of genes with similar expression can explain the diversity of gene expression profiles across spathe developmental stages and colors. A large number of transcripts were assigned to a module with low assignment scores (grey, n Genes = 10,998) indicative of genes that cannot follow a characteristic expression profile. Gene clusters varied in size from 557 (light cyan) to 8774 (turquoise). The large number of genes present in a moderate number of clusters, for such a divergent set of plants, suggested that changes in color across developmental stage was a tightly controlled process.

**Figure 8.**
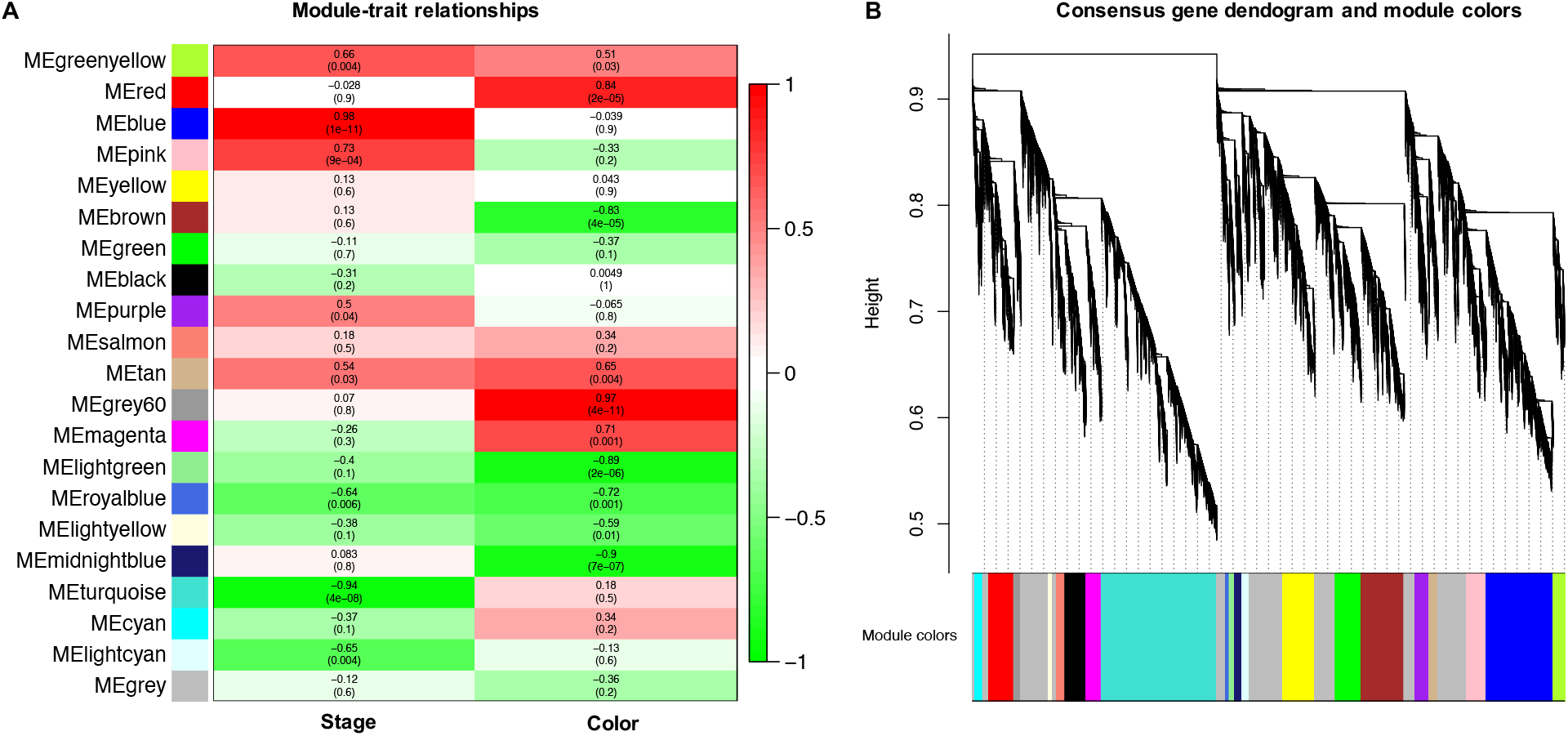
Weighted correlation network analysis of expression across all samples. A. module-trait correlations and corresponding *p*-values (in parentheses). Each row corresponds to a module eigengene (ME), and each column to a trait. The color scale on the right shows module-trait correlations from − 1 (green) to + 1 (red) and, B. Hierarchical cluster tree showing 21 modules of co-expressed genes. Each of the 8,324 DEGs is represented by a leaf in the tree, and each of the modules by a major tree branch. The lower panel shows modules in designated colors.

We focused our attention on the red cluster containing the *CHS* gene (Unigene109494) and used it to identify MYB-like domain proteins that showed similar patterns of expression (Supplemental Table S3). Our analyses showed that there are nine genes encoding MYB domain proteins that showed a highly significantly correlated transcriptional profile compared to *CHS*. Out of these, three MYB encoding genes putatively produced proteins larger than 300 aa, and one of them (Unigene51619) has a target regulatory sequence for miRNA recognition.

## DISCUSSION

In order to gain insight into the molecular events that regulate the spathe color in *A. andraeanum*, we used RNA-seq. This technology provides the opportunity to study the molecular mechanism especially in species where no reference genome is available. Several recent studies have exploited this technology to study traits of particular interest in a wide-range of species due to reduction in library construction and sequencing costs (Garzón-Martínez et al., 2012; Lou et al., 2014; Zhang et al., 2018; Thomas et al., 2019). In particular, it has been applied in *Anthurium* to discover the expression of genes under cold stress (Tian et al., 2013), to identify expression differences between spathes of a red-spathed cultivar and its anthocyanin-loss mutant (Li et al., 2015), to examine expression differences between leaves and spathes of leaf color mutants (Yang et al., 2015) and finally to identify a anthocyanin related MYB transcription factor (Li et al., 2016). Although efforts have been made to elucidate the anthocyanin synthesis pathway in *Anthurium*, developmental stages using different varieties were not specifically examined in any of the previous studies. Therefore, in the present study, we examined the expression patterns of various structural genes (*CHS, F3’H, DFR*, and *ANS*) and TFs (MYB) to determine differences in color in plants with white, red and purple spathes over different developmental stages.

The large diversity in anthocyanin pigmentation in flowers has been an important feature in the coevolution of plants and pollinators. Specifically, for *A. andraeanum*, it has been already established that the spathes accumulate anthocyanins progressively, reaching large quantities at stage 3 and above, and producing other flavonoids (Williams et al., 1981; Spelt et al., 2000). More recent work performed in dark red, red, light red, white, orange and coral varieties of *Anthurium*, which have focused on genes *CHS, F3H, DFR, ANS* and *F3’H*, suggested that *F3’H* expression might be a key control point in the regulation of anthocyanin biosynthesis (Gopaulchan et al., 2014). Additional studies in pink spathe *Anthurium* with different color intensities have also shown that *F3’H* was strongly associated to the intensity of color in the spathe; more specifically, it was shown that the earlier and higher the expression of *F3’H*, the more intense the spathe color (Gopaulchan et al., 2015). Two anthocyanins, cyanidin 3-rutinoside (red) and pelargonidin 3-rutinoside (orange), have been described that confer color to red and pink spathes in *Anthurium* (Iwata et al., 1979, 1985). It has also been established that the shade intensity in red and pink spathes was determined by both the concentration of cyanidin and ratio of cyanidin to pelargonidin (Iwata et al., 1985). In addition, it has been shown that orange and coral spathes are generated by the accumulation of pelargonidin 3-rutinoside pigments with limited or no cyanidin 3-rutinoside (Iwata et al., 1985). Finally, white spathes lack anthocyanins, but contain flavone 6-*C*-glycoside derivatives at similar levels to those found in colored spathes (Iwata et al., 1985), which suggests that the mechanisms regulating differences in the ratio of white to color was downstream in the regulatory cascade in anthocyanin biosynthesis. In concordance, high levels of *CHS* and *F3H* genes during early stages of development has already been established in plants that generate color (Collette et al., 2004). Although previous analyses have implicated *CHS* among other enzymes in the biosynthetic pathway in the determination of color in spathe in *Anthurium*, none have been conclusive and have examined the changes in gene expression using a global approach like the one implemented in this work.

In this study, differences in gene expression during spathe development were observed in the three cultivars. Overall, the *CHS* was the main regulator differentially down-regulated in the white spathe when compared to the red and purple spathes. Besides, the *CHS* in red and purple present similar expression due to is high in early stages and then in late stages its expression decay. This could be due to *CHS* catalyzes the first reaction in the anthocyanin biosynthesis and then in late stages is no need to produce this compound in higher proportion compared to early stages. Similar results related to the high *CHS* content in early developmental stage in different species have been reported previously (Deng et al., 2014; Zhang et al., 2019). However, is different to that reported in petunia flowers where the expression of *CHS* was higher as development stage increased (Sun et al., 2015). In summary, these results confirmed that *CHS* plays an essential role in the biosynthesis of flavonoid in *A. andraeanum* and in the differentiation between white and coloured spathes. In contrast, for the late stage in the purple variety, the color at this stage may also be due to the higher expression levels of *F3’H* than of the other structural genes. Additionally, findings from other studies indicate that genes related with the chalcone isomerase (*CHI)* are coordinately expressed with *F3’H* in the anthocyanin biosynthetic pathway and are, with increased transcript levels toward plant maturity (Ravaglia et al., 2013), for this reason it’s possible that in the case of *A. andraeanum* it is a coordinated expression between *CHS* and *F3’H* to explained the color differences.

Additionally, we detected significant differential expression of a *DFR* gene between white and red, except the direction of the change was opposite to that found in the in other genes, that is that there was an increased expression of *DFR* in the white spathe when compared to the red spathe in the late development stage. This result suggested, that the expression of *DFR* did not contribute to the color differences between the white and colored spathes in this study. Similar findings have also been noted by Gopaulchan et al. (2014), where the white spathes displayed equivalent levels of *DFR* transcript to the red and orange spathes at stage 2, and had higher expression at stage 6 compared the colored spathes. Furthermore, in the purple spathe we identified a cytochrome *P450* oxidase differentially up-regulated when compared to red colored spathe which appeared to determine the difference between the red and purple hues. Functional annotation analysis suggested that the cytochrome *P450* may encode an additional *F3’H* or *F3’5’H*. This latter result was striking because initially the purple spathe displays a red hue at the early stage of spathe development and then acquires the purple hue as it develops into a mature spathe progresses and this cytochrome *P450* could be a highly divergent *F3’H* or *F3’5’H*. Hence, *CHS* may have a stronger influence on anthocyanin accumulation than other structural genes, at least in the early stages of ‘Honduras’ and ‘Rapido’ cultivars. Taken together, our results suggest that *F3’H* is more likely to be a key structural gene in late stages of ‘Purple’ cultivar. Besides, the accumulation of higher amounts of anthocyanin in ‘Purple’ may also be due to a cumulative effect of the high expression levels of all the other structural genes like *P450*. Previous studies in other species indicated that a single gene is not responsible for anthocyanin accumulation and that anthocyanin biosynthesis involves the coordinated mechanism of many genes (Walker et al., 2007; Qian et al., 2014). However, additional research is necessary to test this hypothesis in *A. andraeanum*.

Analyses in different plant systems have focused on understanding the pattern of expression of specific genes and demonstrated how diverse these underlying changes can manifest in differences in coloration can be (Ortiz-Barrientos, 2013; Liu et al., 2017; Ohmiya, 2018). The importance of R2R3 MYB transcription factors in the co-regulation of the anthocyanin biosynthetic pathway have been demonstrated in other species. For example, in *Paeonia ostia*, a study found that two transcription factors, PoMYB2 and PoSPL1, seem to negatively regulate anthocyanin accumulation by interfering with the formation of the MYB-bHLH-WDR complex (Gao et al., 2016). Besides, analyses in *Petunia hybrida* have shown that two bHLH transcription factors interact with a MYB and WD repeat protein to regulate the expression of *CHS* and *DFR* genes (Quattrocchio et al., 1993; de Vetten et al., 1997; Quattrocchio et al., 1999; Spelt et al., 2000; Quattrocchio et al., 2006). In our study, the use of WGCNA to identify co-expression patterns among the six different tissues was especially key to identifying potential regulatory elements for the pathways of interest. Additionally, the mRNA encoding the putative MYB domain contained a silencing RNA-target motif that was polymorphic among samples of white and colored spathes.

## SUPPLEMENTAL MATERIAL

Table S1 contains the differential expressed genes between developmental stages and spathe color. Table S2 contains the functional annotation of DEGs a summary of the statistics for the sequenced data per individual. The clustering according to module colors is provided in Table S3. Figure S1 exhibits the distribution of reads among samples. Figure S2 shows the hierarchical clustering analysis of DEGs.

## Conflict of interests

The authors declare that they have no conflict of interests.

## Data availability

The transcriptome data set supporting the results of this article is available through NCBI ES####### (the corresponding accession numbers will be ready for publication). In addition, data sets supporting the results of this article are included as additional files. Code will be made available through a github repository

## Authors contributions

PU, DG, AML and OEC conceived and designed the experiments. DG collected the samples and performed the RNA isolation experiment. OEC analyzed the data. JAOG helped to analyze the data and draft the manuscript. CQ help in the RNA library construction. PU, DG and OEC coordinated the study and revised the manuscript. All authors read and approved the final manuscript.

## Acknowledgments

The authors are very grateful to Jennifer Avey and her staff at Kairi Blooms Ltd. Trinidad, for providing the plant material used in this study.

## Abbreviations

ANS: anthocyanidin synthase
bHLH: basic helix-loop-helix
CHS: chalcone synthase
DFR: Dihydroflavonol 4-reductase
DEG: differentially expressed gene
FDR: false discovery rate
F3’H: flavonoid 3’-hydroxylase
F3’5’H: flavonoid 3’5’ hydroxylase
GO: gene ontology
MDS: multidimensional scaling
UPGMA: unweighted pair group method with arithmetic mean
RNA-seq: RNA sequencing
TPM: total number of transcripts per million
WGCNA: weighted correlation network analysis

## Notes

### Competing Interest Statement

The authors have declared no competing interest.

